# Spatial and Spectral Components of the BOLD Global Signal in Rat Resting-State Functional MRI

**DOI:** 10.1101/2022.12.06.519194

**Authors:** Nmachi Anumba, Eric Maltbie, Wen-Ju Pan, Theodore J. LaGrow, Nan Xu, Shella Keilholz

**Affiliations:** Department of Biomedical Engineering at Georgia Institute of Technology and Emory University; School of Electrical and Computer Engineering at Georgia Institute of Technology

## Abstract

In resting-state fMRI (rs-fMRI), the global signal average captures widespread fluctuations related to unwanted sources of variance such as motion and respiration, and has long been used as a regressor to reduce noise during data preprocessing. However, coherent neural activity in spatially-extended functional networks can also contribute to the global signal. The relative contributions of neural and non-neural sources to the global signal remain poorly understood. This study sought to tackle this problem through the comparison of the blood oxygenation level dependent (BOLD) global signal to an adjacent non-brain tissue signal from the same scan in rs-fMRI obtained from anesthetized rats. In this dataset motion was minimal and ventilation was phase-locked to image acquisition to minimize respiratory fluctuations. In addition to contrasting the spatial and spectral components of these two signals, we also observed these differences across the use of three different anesthetics: isoflurane, dexmedetomidine, and a combination of dexmedetomidine and light isoflurane. Here, we report differences in the spectral composition of the two signals as evaluated by a power spectral density (PSD) estimate and a fractional amplitude of low-frequency fluctuations (fALFF) calculation. Additionally, we show spatial selectivity for specific brain structures that show an increased correlation to the global signal both statically and dynamically, through Pearson’s correlation and co-activation pattern analysis, respectively. All of the observed differences between the BOLD global signal and the adjacent non-brain tissue signal were maintained across all three anesthetic conditions, showing that the global signal is distinct from the noise contained in the tissue signal. This study provides a unique perspective to the contents of the global signal and their origins.

## INTRODUCTION

In resting-state functional magnetic resonance imaging (rs-fMRI), there is a unique opportunity to study the brain’s intrinsic functional organization as captured by low-frequency fluctuations in the blood oxygen level dependent (BOLD) signal (Biswal et al., 1995). However, the spontaneous BOLD fluctuations are small (Pais-Roldán et al., 2018) and contributions arising from neural activity can easily be swamped by signal changes related to motion and respiration (Murphy et al., 2013; Power et al., 2017). To minimize unwanted signal changes from non-neural sources, the global average of the BOLD signal from all of the voxels in the brain (‘global signal’) is often regressed from the data during preprocessing for both task-based fMRI and rs-fMRI data (Aguirre et al., 1998; Macey et al., 2004; Murphy & Fox, 2017; Zarahn et al., 1997). Despite the success of global signal removal as a denoising step (Birn et al., 2006; Parkes et al., 2018; Satterthwaite et al., 2012; Wise et al., 2004; Yan et al., 2013), regression of the signal remains a controversial practice as many argue that the global signal contains neural data of interest (Ciric et al., 2017; Fox et al., 2009; T. T. Liu et al., 2017; Murphy & Fox, 2017; Saad et al., 2012).

The removal of the global signal from rs-fMRI data is primarily motivated by the principle that the majority of widespread fluctuations that are captured by this global average are likely to be the result of widespread sources of noise. Major sources of noise that are known to be reflected in the global signal are motion (Parkes et al., 2018; Satterthwaite et al., 2012; Yan et al., 2013) and physiological factors such as cardiac pulsation, vascular tone, and changes in cerebral blood flow (Birn et al., 2006; Wise et al., 2004). These types of noise tend to be reflected in large portions of the brain signal, even at times in the whole image, and their influence on image quality can be significantly reduced through removal of the global signal (Ciric et al., 2017; Fox et al., 2009; T. T. Liu et al., 2017; Murphy & Fox, 2017; Saad et al., 2012). These types of noise affect both the brain signal and that of anatomical structures surrounding the brain. Imaged features such as the skull or surrounding head tissue would also be affected by head motion, any whole-image scanner noise, and magnetic field shifts as a result of respiratory or other forms of motion. When comparing human and rodent imaging, the imaged head tissue in humans is minimal, whereas there is substantially more tissue surrounding the brain in rodents. Therefore, a rodent study comparing the global signal to the signal contained in these surrounding structures has the potential to elucidate the contributions of specific noise to the global signal.

The data used in this study was acquired using a protocol that was optimized to minimize motion and respiratory noise to better isolate the non-noise components that are captured in the BOLD signal. These noise-limiting steps included mechanical ventilation, the use of a paralytic, employment of stereotaxic head fixation, and respiratory phase-locked image acquisition, which minimizes respiratory contributions to the signal. With these procedures in place, we aimed to study the global signal in the context of minimal noise. An important tool for minimizing noise in rodent rs-fMRI is the use of anesthesia, which allows animals to be head-fixed without pain or stress and prevents voluntary motion that can affect the magnetic field. However, the anesthetic agents used can affect neurovascular coupling (Franceschini et al., 2010) and induce potential confounds into the rs-fMRI data. Two of the most commonly used anesthetics for rodent rs-fMRI include isoflurane and dexmedetomidine. Each of these anesthetic agents can induce changes in vascular tone and cerebral blood flow. Isoflurane is a vasodilator which increases cerebral blood flow (Sicard et al., 2003) while dexmedetomidine is a vasoconstrictor which decreases cerebral blood flow (Ganjoo et al., 1998). These effects can influence measurements of functional connectivity (Grandjean et al., 2014) in rs-fMRI and even alter which frequency bands exhibit the greatest coherence between rs-fMRI and local field potentials (Pan et al., 2013). A common method in rodent rs-fMRI is to use a combination of both isoflurane and dexmedetomidine together because the vasodilatory effects of isoflurane have been shown to be attenuated by the opposing vascular effects of dexmedetomidine (Ohata et al., 1999).

In this work, we investigated the spatial and spectral differences between the brain global signal and a same-scan adjacent non-brain tissue signal across three different anesthetic conditions: (1) isoflurane, (2) dexmedetomidine, and (3) a combination of the two (isodex). The use of multiple anesthetics ensures that results are not specific to a single anesthetic condition. We examined how the tissue and brain signals differed through a power spectral density analysis, a voxel-wise correlation analysis, and a dynamic co-activation pattern (CAP) analysis. We provide evidence that the global signal contains specific spectral and spatial qualities that are separate from noise and could be linked to brain activity. Additionally, these findings were robust across the use of the three different anesthetic protocols. The results of this study provide unique insight into the composition of the global signal.

## METHODS

### Animal preparation

All protocols were approved by the Institutional Animal Care and Use Committee (IACUC) at Emory University (Atlanta, GA, USA) and all procedures were performed in strict compliance with the IACUC protocols. Data was acquired from eight male Sprague Dawley rats (299g – 339g, Charles River). Animals were inducted under 5% isoflurane and were maintained at 2% isoflurane during handling and placement in the MRI cradle. All rats were intubated and underwent ventilator-assisted breathing at a rate of 1 Hz. An infusion line was inserted subcutaneously to administer the paralytic pancuronium at a rate of 1.5 mg/kg/hr for the duration of the scan. Animals were placed in a homemade custom MRI cradle and positioned with their teeth secured in a bite bar and their heads fixed using ear bars.

After preparation was complete, the animals underwent a scanning session in which scans were consecutively collected under three different anesthetic protocols: isoflurane alone, dexmedetomidine alone, and a combination of dexmedetomidine with light isoflurane (isodex). Initially, animals were scanned under 2% isoflurane as part of a separate study; however these scans were not included in the analyses for this paper as deep levels of isoflurane have been shown to result in burst suppression and increase the spatial extent of contribution to the global signal (X. Liu et al., 2013). Afterwards, isoflurane was reduced to 1.5% and the animals were scanned for an average of 35 minutes. After the isoflurane scan, the animals were subcutaneously injected with a 0.025 mg/kg bolus of dexmedetomidine, taken off isoflurane five minutes later, and then switched to a 0.05 mg/kg/hr subcutaneous infusion of dexmedetomidine at 10 minutes post-bolus. The animals were then scanned again while exclusively under dexmedetomidine for an average of about 61 minutes. Functional dexmedetomidine scans were started an average of about 14 minutes after isoflurane was disconnected. Afterwards, the animals were introduced to a low dose of isoflurane at 0.5% in combination with the 0.05 mg/kg/hr subcutaneous infusion of dexmedetomidine. The animals were then scanned for a final time while under dexmedetomidine and low isoflurane (isodex) for about 49 minutes on average. Functional isodex scans were started an average of 17 minutes after the low dose of isoflurane was introduced. A table with the exact timing of anesthetic introduction and image acquisition for each rat is included in the supplementary section (**Supplementary Table 1**). In order to use the maximum amount of data possible in accordance with the length of the shortest scan, only the first 30 minutes of the isoflurane and dexmedetomidine scans and the first 20 minutes of the isodex scans were used for the group analysis. The last 30 minutes of dexmedetomidine scans were also analyzed to compare for any unwanted effects of lingering isoflurane that could be present in the first 30 minutes of these scans; these findings are reported in the supplemental section (**Supplementary Figure 1**). Physiological parameters of the animal were measured using a pulse oximeter that was placed on a hindpaw of the animal and through a rectal temperature probe. Temperature of the animal was maintained around 37 ± 0.5 °C using a heated water bath system.

### Data acquisition

MRI data was acquired using a 20 cm horizontal bore 9.4 T Bruker Biospec MRI and a homemade transmit-receiver surface coil. A T2-weighted RARE anatomical scan was acquired for each rat (TR = 3500 ms, TE = 11 ms, 24 axial slices, 0.5 mm3 isotropic voxels). All functional rs-fMRI scans were acquired using a gradient-echo echo-planar imaging (EPI) sequence with the following parameters: partial Fourier encoding with a factor of 1.4, field of view (FOV) of 35×35 mm2, matrix size of 70×70, isotropic voxel size of 0.5 mm^3^, 24 axial slices for whole-brain coverage, a flip angle of 68.4°, bandwidth = 216.45 kHz, TE = 15 ms, and TR = 2000 ms. A 3-volume reversed blip EPI image with the aforementioned parameters was acquired before each longer functional scan for topup correction (Andersson et al., 2003; Smith et al., 2004). All functional EPI scans included saturation bands to reduce signal from non-brain structures and were preceded by 10 dummy scans.

All functional EPI scans for this study were phase-locked, meaning that the acquisition of the these images was set to a multiple of the animal’s respiratory rate so that each image was acquired during the same phase of the respiratory cycle (Pan et al., 2020). This practice was adopted here to limit any effects of motion that could arise due to movement of the chest cavity and volumes being imaged at different points in the respiratory cycle. In the case of this study, images were acquired at a frequency of every other breath, or 2 Hz (TR = 2000 ms).

### Preprocessing

All preprocessing of the data used in this study was performed using the following software: FSL (Jenkinson et al., 2012), Analysis of Functional NeuroImages (AFNI) (Cox, 1996), and ITK-SNAP (Yushkevich et al., 2006). Distortion correction was applied to all scans using FSL Topup and volume registration to the 30th volume of each scan was performed using AFNI 3dVolReg. Motion regression (6 parameters and up to 2 polynomials) and bandpass filtering (0.01 – 0.2 Hz) were performed in one step using AFNI 3dTProject. The frequency filter range was chosen to compare all anesthetic conditions across the same frequencies, according to previous work that has shown contributions from higher frequencies at resting state under dexmedetomidine (0.01 – 0.25 Hz) than under isoflurane (0.01 – 0.1 Hz) (Pan et al., 2013). Spatial smoothing was not applied to this data as smoothing the non-brain tissue signal may have had differential effects due to its size being much smaller than the brain. Brain masks for each scan were acquired automatically in AFNI using 3dAutomask. Individual non-brain tissue masks were acquired manually using ITK-Snap and created to consist of 500 – 600 voxels for each scan (**Figure 1**). Areas of tissue closest to the brain were chosen for these masks in an effort to obtain good signal and maximize similarities in the types of noise that would affect both signals. The individual brain and non-brain tissue masks were then used to segment the two signals for analysis. For group average signals, all scans were aligned to a single subject using direct, linear EPI to EPI registration via AFNI 3dAllineate. For the group voxel-wise correlation analysis, scan segments, global signals, and non-brain tissue signals from each rat were concatenated.

**Figure 1:**
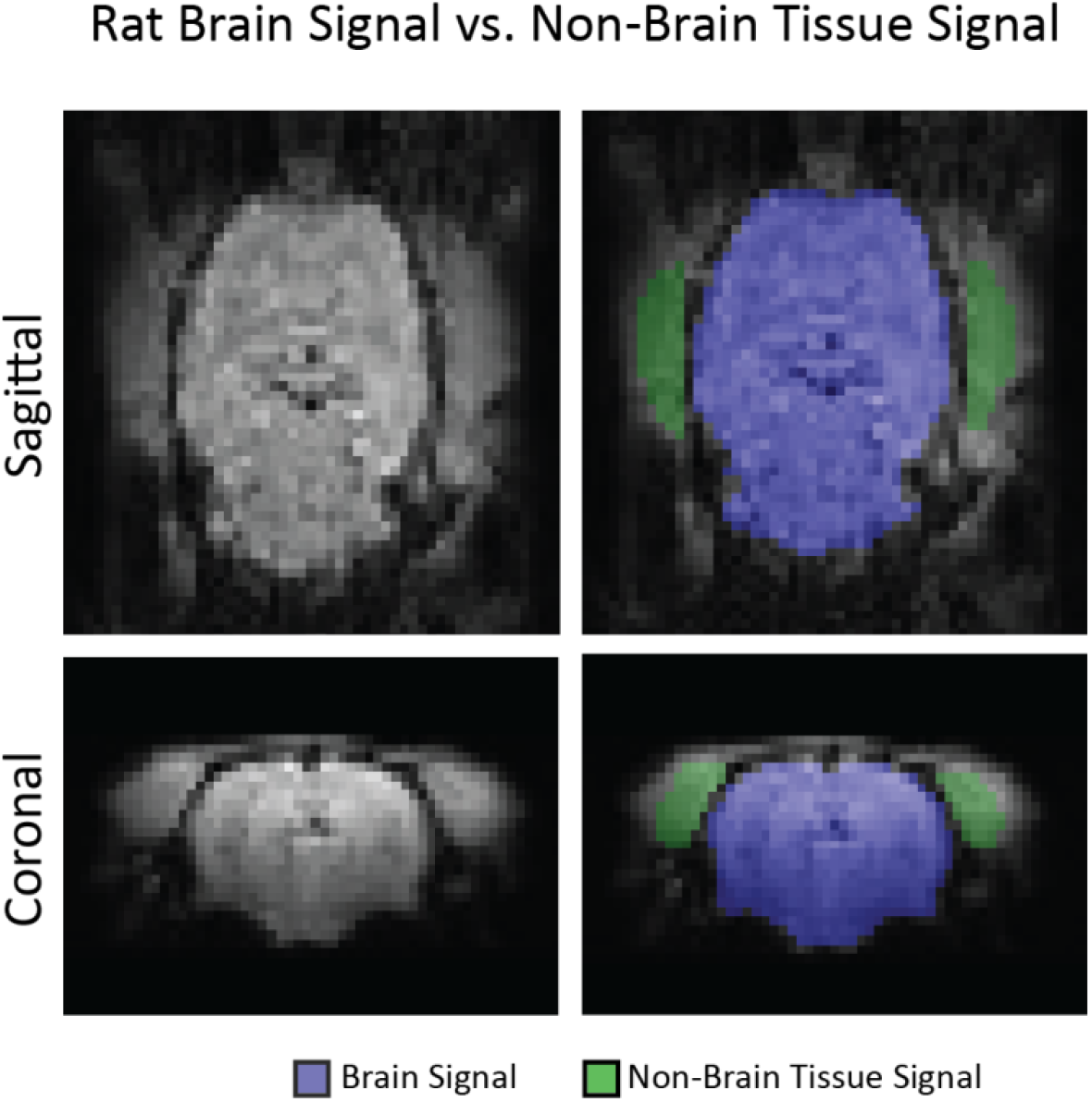
Rat Brain Signal vs. Non-Brain Tissue Signal. Rat anatomy results in notably more non-brain tissue being imaged around the brain than is observed in human imaging. This figure shows slices from a functional scan of subject 6 (left column) and then shows those same slices with the actual non-brain tissue mask used in this study as well as a representative brain mask (right column). The substantial amount of tissue imaged alongside the brain in rats allowed us to carry out these adjacent signal comparisons.

### Global signal comparison to non-brain tissue signal

A comparison was made between the global signal and the averaged signal of the surrounding non-brain tissue in the same scan. This was done to identify any characteristics of the global signal that could be consistently linked to brain activity and were separate from noise. The rationale between comparing these two signals comes from the fact that despite various measures being taken to minimize noise during acquisition, both the global signal and the non-brain tissue signal would be affected by any residual motion, scanner noise, and magnetic field shifts that were to occur. Therefore, any signal attributes that are the result of brain activity would stay confined within the global signal, whereas any signal characteristics that primarily reflect noise would potentially be observed in both signals. In this way, the non-brain tissue signal acts as a comparable noise measure.

All analyses of these signals were performed in MATLAB using custom scripts. The global signal was calculated by averaging the timecourses of all voxels within the brain and then z-scoring the signal by subtracting its mean and dividing by its standard deviation. The non-brain tissue signals were calculated in a similar manner, by averaging the timecourses of all voxels within the segmented tissue region and z-scoring them.

### Power analysis

Power spectral density (PSD) estimates of the global signal were compared to those of the non-brain tissue signal to evaluate the various frequency contributions within the range of 0.01 – 0.2 Hz. PSDs were computed in MATLAB using Welch’s method with a Hamming window and 50% overlap. The power spectral densities found for each individual scan were averaged to create the group average for each anesthetic group. These power spectral densities were then normalized to a scale of 0 to 1 in order to compare the power distributions of the two signals. The fractional amplitude of low-frequency fluctuations (fALFF), as defined by Zou *et al. (Zou et al*., *2008)*, was calculated for each power spectrum by summing the square root of the power within the 0.01 – 0.08 Hz range and dividing this by the sum of the square root of the power across the entire frequency range (0.01 – 0.2 Hz). A voxel-wise power distribution map was calculated by summing the power at each voxel over a specified frequency range and displaying this number in decibels. The frequency ranges used were 0.01 – 0.1 Hz for isoflurane scans and 0.01 – 0.25 Hz for dexmedetomidine and isodex scans.

### Signal spatial correlation analysis

The global signal and the non-brain tissue signal for each scan were correlated using a Pearson’s linear correlation. A voxel-wise correlation analysis was also performed in which a Pearson’s linear correlation was calculated between the global signal and each voxel timecourse within the brain, and again with the non-brain tissue signal and each voxel timecourse within the brain.

### Co-activation pattern analysis

In order to compare the non-brain tissue and brain global signals as they vary over time, as opposed to the time-averaged comparison obtained from correlation, we performed a co-activation pattern (CAP) analysis. CAP analysis is a dynamic analysis method that was first proposed by Liu and Dyun (X. Liu & Duyn, 2013). This method involves choosing a seed region or region of interest (ROI) and setting a threshold for that ROI signal. The time points at which the ROI signal passes the threshold are used to extract fMRI volumes, which are then clustered and averaged to produce different CAPs. In this study, two separate CAPs were calculated by using the global signal and the non-brain tissue signal as distinct ROIs. These two CAPs were calculated individually for each scan and the group CAPs are the result of an equal average of the individual CAPs for that group. The ROI signal timecourses were z-scored, and the threshold was set to a z-score greater than 1. For each ROI, all frames at time points that exceeded the threshold were averaged resulting in the global signal and non-brain tissue signal CAPs.

## RESULTS

We studied both spectral and spatial features of the global signal in eight male Sprague Dawley rats. These characteristics were compared to a same-scan non-brain tissue signal that was segmented and compared to the global signal of each rat. The individual analyses of each rat can be found in the supplemental section (**Supplemental Figures 2-9**), whereas group results are displayed in this section.

### Power distribution results

We created a PSD estimate for both the global signal and the non-brain tissue signal for each scan (**Figure 2A**). The group average power spectra are shown for scans under isoflurane, dexmedetomidine, and isodex. The comparison of the contributing frequencies in both signals under the various conditions shows distinct differences between the global signal and the non-brain tissue signal. Most notably, the tissue signal appears to exhibit a higher proportion of low-frequency contribution to the overall signal. To evaluate this, we calculated the fractional amplitude of low-frequency fluctuations (fALFF) (Zou et al., 2008) for each trace. The fALFF values for the global signal were 0.4398, 0.4652, and 0.4393 for isoflurane, dexmedetomidine, and isodex, respectively. However, across all anesthetic conditions an increase in the fALFF value is observed for the non-brain tissue signal with fALFF values being 0.5181, 0.4811, and 0.5049 for isoflurane, dexmedetomidine, and isodex, respectively. The differences displayed in these power spectral plots emphasize the distinct spectral constitutions of these two signals, implying that there may be certain activity captured in the global signal that is not being captured in the non-brain tissue signal.

**Figure 2:**
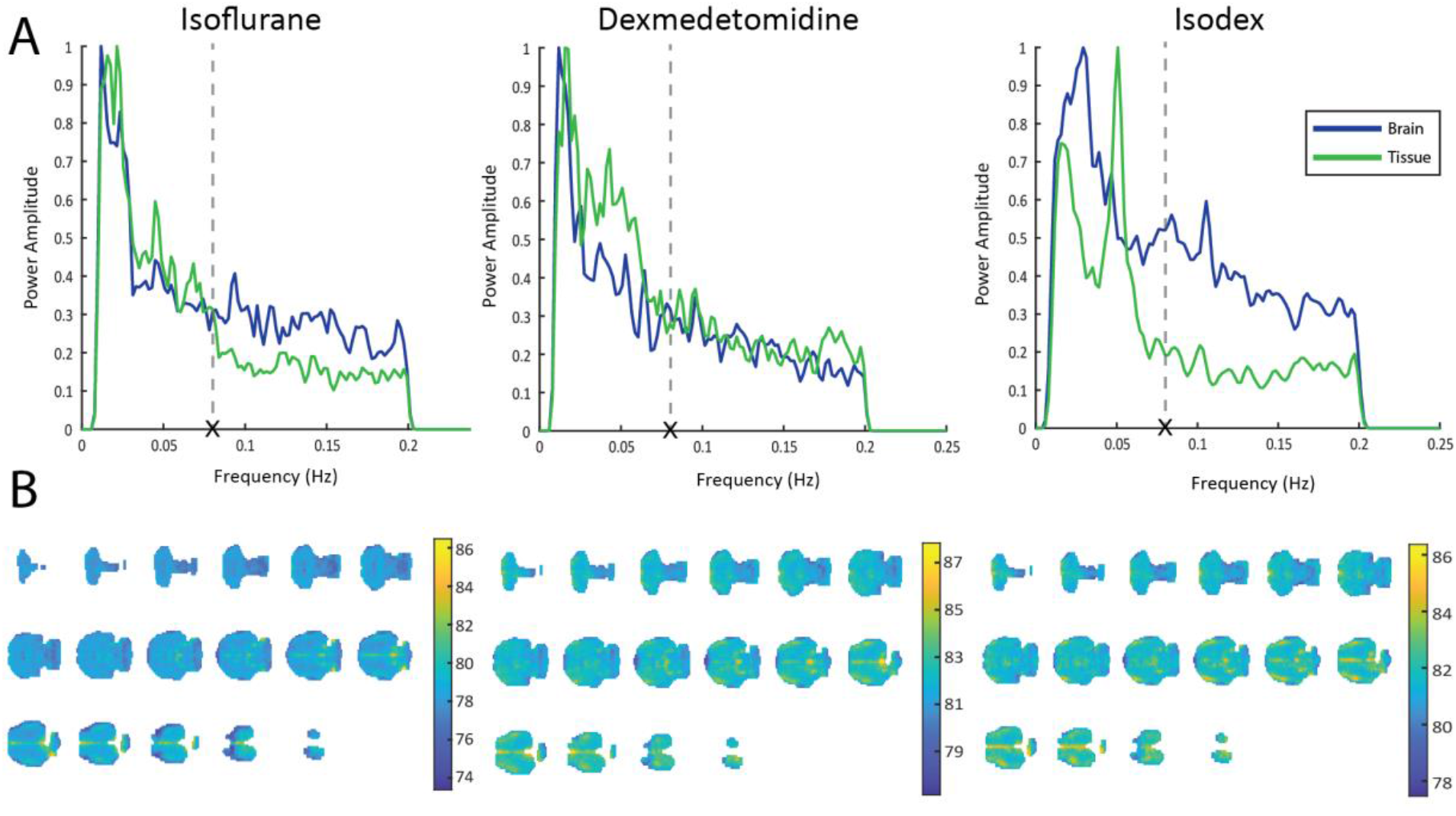
Power distribution of the brain global signal and the tissue signal. (A) Power spectral density estimates for the global signal (blue) and the non-brain tissue signal (green) across the three anesthetic conditions. The X and dashed line mark 0.08 Hz as used for the fALFF calculation. Under all three anesthetic conditions the tissue signal had a higher fALFF value than the brain global signal. (B) Group level spatial distribution of power across the brain under all anesthetic conditions. High levels of power were localized to voxels along the posterior midline of the brain and, in the case of dexmedetomidine and isodex, bilaterally in the cortex.

In addition to this, we observed the power distribution of the brain signal across all three anesthetic conditions (**Figure 2B**). For scans acquired under isoflurane, we summed the power for each voxel over the 0.01 – 0.1 Hz range, whereas for dexmedetomidine and isodex scans we summed the power over the 0.01 – 0.25 Hz range. Results are reported in decibels. Across all three anesthetic conditions the highest amount of power is observed in a posterior medial part of the brain and throughout the entire medial section in the more superior slices of the brain.

### Non-brain tissue vs. brain results

In order to show the distinct temporal nature of the global signal from the non-brain tissue signal, we performed a Pearson’s linear correlation to correlate the two signals for each scan at an individual level (**Figure 3**). The mean values of correlation from all eight rats in each of the anesthetic conditions were -0.0038, 0.0072, and 0.0313 for isoflurane, dexmedetomidine, and isodex, respectively. When analyzed with a one-sample t-test, these mean values were not statistically different from zero (isoflurane: p = 0.8943; dexmedetomidine: p = 0.7925; and isodex: p = 0.1357). The correlation values for all rats across anesthetic conditions were within the range of 0.15 to -0.15, reflecting little to no correlation. The lack of correlation between the global signal and the non-brain tissue signal show that even on an individual rat level there is very little similarity between these two signals and provides evidence that they contain different information.

**Figure 3:**
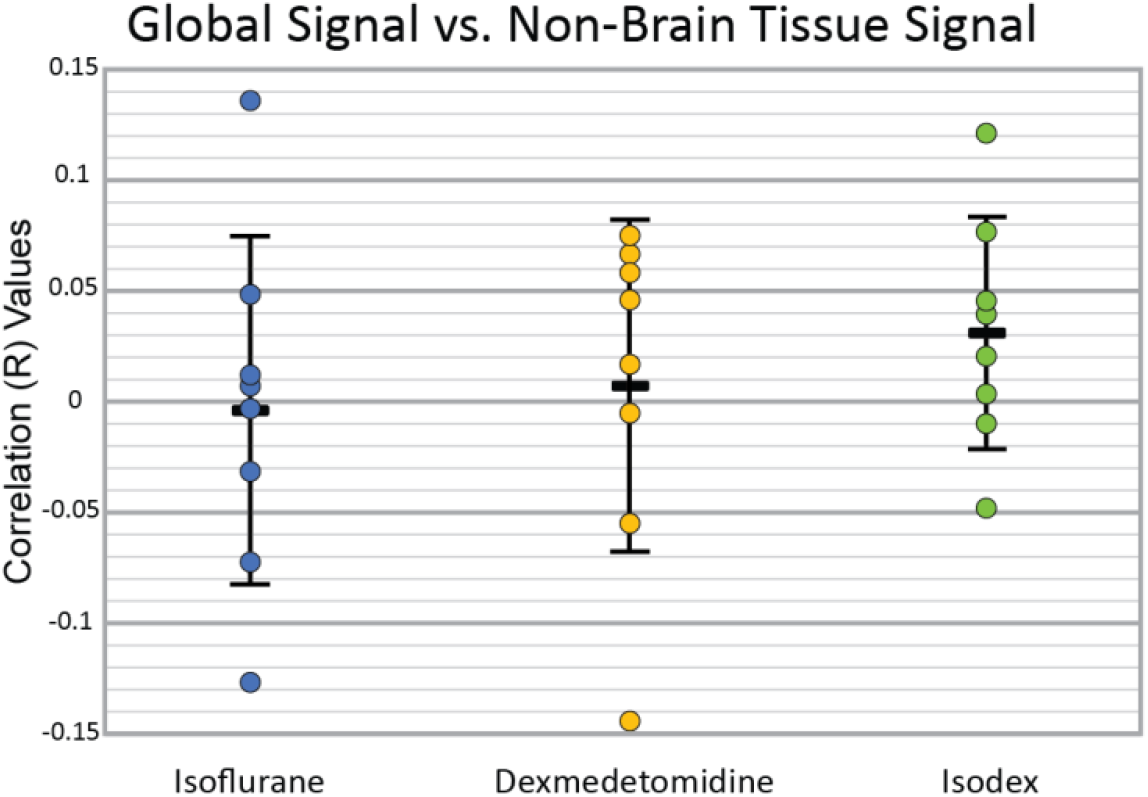
Correlation values between the brain global signal and the non-brain tissue signal. The correlation between the global signal and the non-brain tissue signal for each rat across anesthetic conditions. Black rectangles represent the means for each group: isoflurane (blue) = -0.0038, dexmedetomidine (yellow) = 0.0072, isodex (green) = 0.0313. These means were not statistically different from zero (p = 0.8943, 0.7925, and 0.1357, respectively). The low R values indicate very little correlation between the two signals.

In order to demonstrate the spatial distribution of the global signal, we performed a voxel-wise correlation analysis (**Figure 4**). A Pearson’s correlation between the global signal and the timecourse of each voxel within the brain mask was computed. Similarly, we also computed the correlation between the non-brain tissue signal and the timecourse of each voxel. This comparison was made to display the fact that spatial patterns observed with the global signal were not the result of whole-image noise. The spatial correlation to the global signal across all anesthetic conditions showed higher correlation in an anterior medial part of the brain. In isoflurane scans, higher correlation is also shown bilaterally and throughout the outer cortical regions. Correlation to the global signal in dexmedetomidine scans was overall slightly lower and showed a weaker correlation than that seen bilaterally in cortex under the other two conditions. In isodex scans, there was also a high bilateral correlation to the global signal in the anterior cortex. When comparing the global signal correlation to the non-brain tissue signal correlation, it can be seen that the tissue signal correlation was overall much lower and that there were no detectable spatial patterns. The individual results of this analysis in each rat can be seen in the supplementary section (**Supplementary Figures 2-9**).

**Figure 4:**
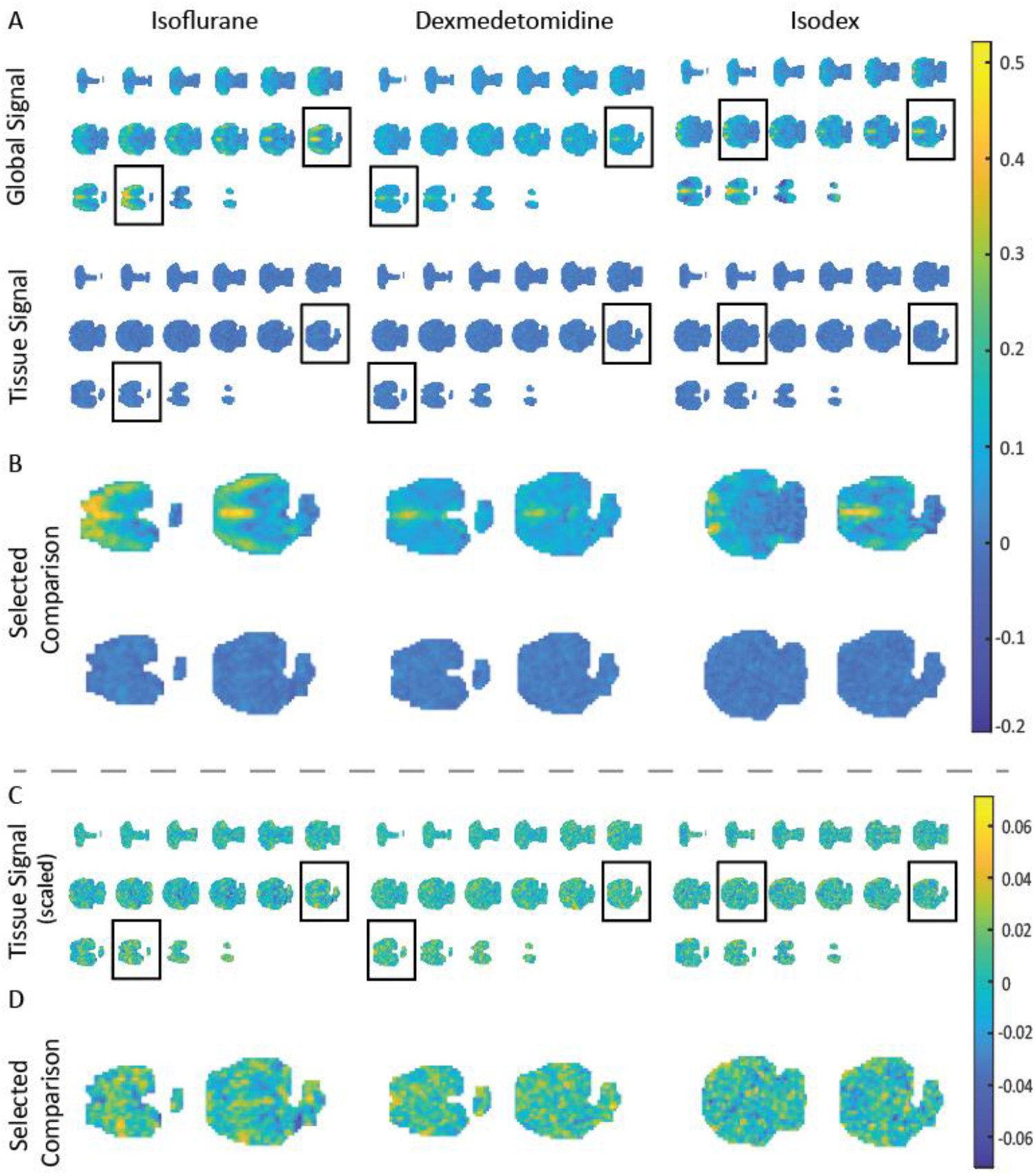
Spatial correlation to the brain global signal and the non-brain tissue signal. (A) The correlation of each voxel to the global signal (top row) and to the tissue signal (bottom row). Specific slices are outlined by a black box and enlarged in the following section for a clearer comparison. High correlation to the global signal is observed in an anterior medial part of the brain across all anesthetic conditions and bilaterally under isoflurane and isodex. Overall correlation to the non-brain tissue signal is much lower. (B) The enlarged slices that were highlighted in the previous section, show a clearer comparison between correlation to the global signal (top row) and to the tissue signal (bottom row). (C) Voxel-wise correlation to the tissue signal with the color bar adjusted so that spatial differences can be observed. The same slices outlined in (A) are highlighted and enlarged in (D) where no spatial specificity in correlation to the tissue signal can be observed.

### CAPs results

To further emphasize the spatial differences between the global signal and the non-brain tissue signal, we performed a co-activation pattern (CAP) analysis (**Figure 5**). In this case, the two “ROIs” used to calculate the CAPs were the global signal and the non-brain tissue signal. Unsurprisingly, the global signal CAPs show high activation from similar brain regions that showed high correlation to the global signal in the voxel-wise analysis (**Figure 4**). In the isoflurane scans, the global signal CAP displays high activation bilaterally throughout cortex, and in some subcortical structures, as well as medially. The dexmedetomidine scans show lower levels of activation in the global signal CAP when compared to the other two conditions, however, higher levels of activation are observed both medially and bilaterally in cortex. The global signal CAP from the isodex scans shows high activation in medial cortex as well as bilaterally in anterior regions of cortex. In comparison, the non-brain tissue signal CAPs across all anesthetic conditions display much lower activation overall and little to no detectable spatial specificity.

**Figure 5:**
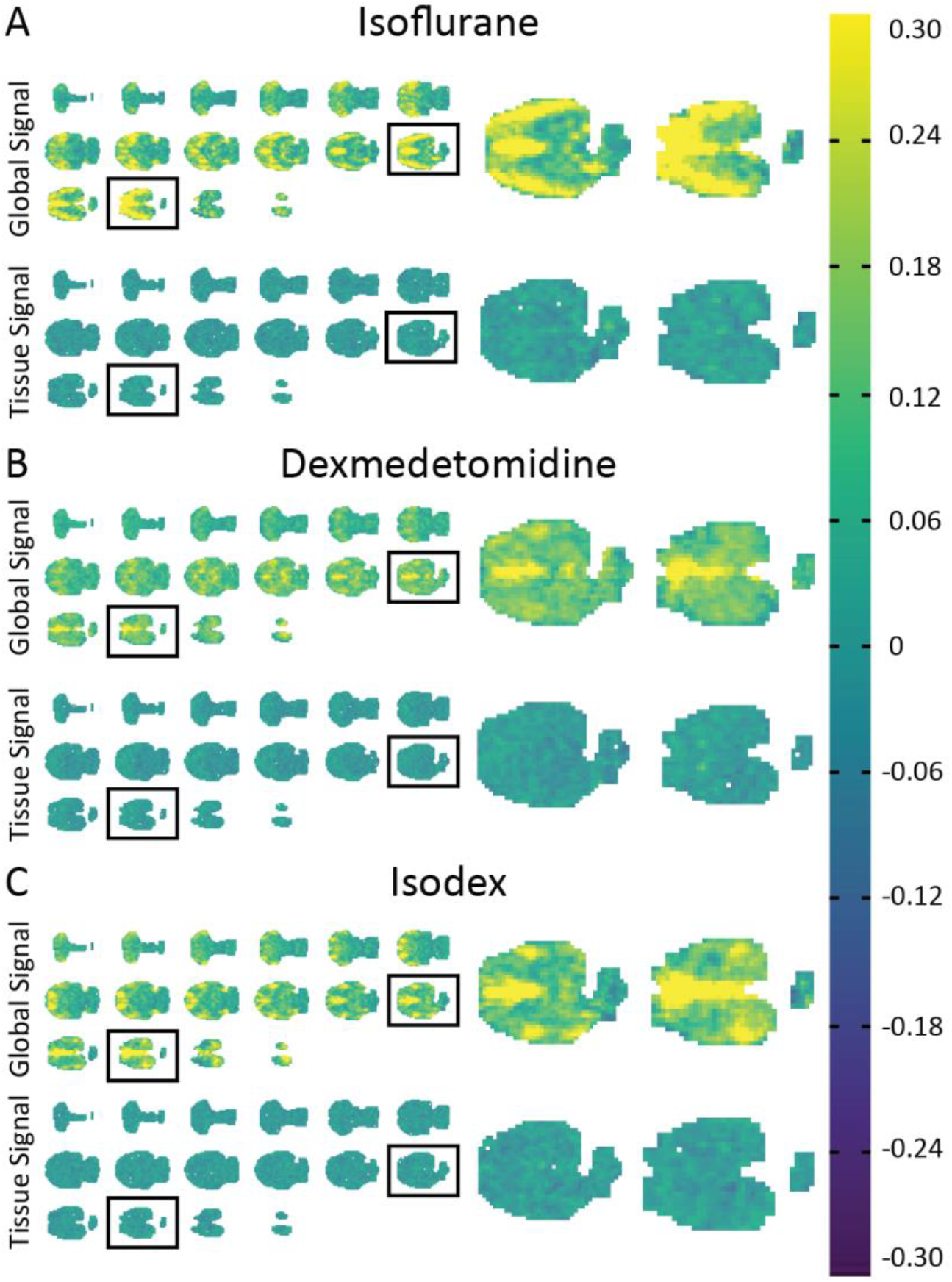
Z-scored co-activation patterns (CAPs) to the global signal and the non-brain tissue signal. The average CAPs for scans acquired under isoflurane (A), dexmedetomidine (B), and isodex (C) when the global signal and the tissue signal were used as regions of interest. The global signal CAP showed clear high activation in certain areas of the brain across all anesthetic conditions. These brain regions are very similar to those that showed high correlation to the global signal in the correlation analysis shown in Figure 4. In contrast, the non-brain tissue signal CAP displays overall lower activation throughout the brain and undetectable spatial specificity.

## DISCUSSION

The use of global signal regression in preprocessing for rs-fMRI remains controversial. While the global signal captures widespread fluctuations related to motion and physiological cycles (Birn et al., 2006; Parkes et al., 2018; Satterthwaite et al., 2012; Wise et al., 2004; Yan et al., 2013), spatially-extended patterns of neural activity are also included (Ciric et al., 2017; Fox et al., 2009; T. T. Liu et al., 2017; Murphy & Fox, 2017; Saad et al., 2012). In an animal model where both motion and physiological noise are minimal, we have shown differences in spatial and spectral distribution between the global signal and a comparable noise signal that are similar under multiple anesthetic conditions. This work provides strong evidence that much of the global signal arises from neural sources in this well-controlled model.

### Potential DMN contribution to the global signal

The correlation to the global signal and its spatial distribution of power appeared to be highly localized along the midline of the brain, regardless of the anesthetic used (**Figure 4**). No such spatial specificity was observed when correlating voxel timecourses to the non-brain tissue signal. Upon further examination of these brain areas, we found that they show striking resemblance to structures that have been identified to form the default mode network (DMN) in rats (Liang et al., 2012; Lu et al., 2012; Upadhyay et al., 2011).

In humans, areas of high correlation to the global signal are most notable along the posterior midline and in the occipital cortex (Billings & Keilholz, 2018; Power et al., 2017). These human studies also noted relatively low correlation in frontal cortex. The reason behind these regional differences between rat and human studies could be largely due to anatomical and functional differences in brain organization across species (Xu et al., 2022). For example, when considering the midline, higher correlation in humans was observed in posterior regions, whereas in rats, higher correlation is observed along the anterior midline. This is in accordance with defined regions that make up that DMN for each species; the precuneus and posterior cingulate in humans (Thomas Yeo et al., 2011) and the cingulate and retrosplenial cortices in rats (Lu et al., 2012). Additionally, the human data contained notably more signal contribution from head motion and respiratory patterns (Power et al., 2017), factors that were largely accounted for in our rat study. The inclusion of these additional signals could also contribute to the regional differences observed between the studies.

The areas of localization of power in this study were similar to those that showed high correlation to the global signal (**Figure 2**). However, the areas that show high power in the rats, presumably reflecting areas with high neural or metabolic activity, also show similarity to the brain areas that show high correlation to the global signal in humans, namely the posterior midline. This discrepancy highlights that there may be some underlying activity in certain structures that results in high power but does not necessarily contribute to the global signal.

*Power et al*., highlight the variability of spatial distribution of the global signal across individuals, scanning protocol used, and hardware artifacts (Power et al., 2017). In contrast, the current study saw similar results on an individual level (**Supplementary Figures 2 – 9**), possibly due to the strict control of motion and respiration. A future study using a different scanner and scanning protocol could be performed to further validate the consistency of these results.

Similar to a previous study done in humans (X. Liu et al., 2018), CAP analysis based on peaks of the global signal identifies similar brain regions to those that show higher correlation to the global signal. Similar to the static correlation results, a non-brain tissue signal CAP showed little to no spatial coherence and lower levels of activation.

The findings of this paper indicate that DMN-like brain areas contribute a disproportionate amount of activity to the global signal in rats. This is significant in that it means that preprocessing steps that eliminate the global signal in rodent studies are most likely removing potentially relevant information from large-scale brain activity, including contributions from large neural networks like the DMN and potentially whole-brain dynamics. However, the contribution of noise to the global signal could be stronger in freely breathing and awake animals.

### Effects of anesthesia on results

In this study, we looked at the differences contained in the global signal and a comparable same-scan non-brain tissue signal. In order to account for differences in brain signal that arise due to the use of an anesthetic (X. Liu et al., 2013; Masamoto et al., 2009; Masamoto & Kanno, 2012), we carried out this analysis in data collected under three different anesthetic protocols. Despite the mechanistic differences across these anesthetics, the results found were consistent regardless of the type of anesthetic used. This is especially apparent in **Figure 4** and **Figure 5**, where similar brain areas are shown to have higher static and dynamic correlation to the global signal under isoflurane, dexmedetomidine, and isodex.

The anesthetic effects of isoflurane are mediated by a complex interaction with GABAergic neurotransmission (Ferron et al., 2009) through a completely different mechanism than the sedative effects of dexmedetomidine which are mediated by interaction with alpha2-adrenergic receptors (Ganjoo et al., 1998). Additionally, isoflurane is a vasodilator which increases cerebral blood flow (Sicard et al., 2003) while dexmedetomidine is a vasoconstrictor which decreases cerebral blood flow (Ganjoo et al., 1998). Hence, why the combination (isodex) is commonly used to produce an optimized anesthetic state (Grandjean et al., 2014). Deeper levels of anesthesia have been shown to cause less spatially-specific functional connectivity across the brain (X. Liu et al., 2013). To minimize disruption to natural brain activity, the anesthetic levels used in this study were kept relatively light, and as a result, we continued to see spatially-specific correlation to the global signal across all three anesthetic conditions.

One advantage of performing a study in humans or awake animals would be the absence of such effects, although this absence also comes with the inclusion of more inherent signal noise. This tradeoff however, does not negate the utility of using anesthetized rodents as they present a unique situation in which to study neural aspects of the rs-fMRI signal (Xu et al., 2022). These findings, in addition to the application of rodent disease models and genetic manipulation tools, are invaluable to our understanding of human brain function.

Despite the anticipated anesthetic-induced effects on brain activity, the results of the present study demonstrate consistent non-random correlation to global brain signal with a similar spatial distribution across anesthetic conditions, indicating that these results are unlikely to be artifacts of the anesthesia.

### Limitations of the study

One limitation of this study is the fact that the timing of introduction for the anesthetics varied from scan to scan, due to differences in time required for setup and maintenance of physiological conditions. Despite this difference, similar results were found across scans when these analyses were done on an individual basis (see **Supplemental Figures**).

### Conclusions

Our findings provide strong evidence that the global signal and the acquired non-brain tissue signal contain different information and that the information contained in the global signal is likely to contain substantial contributions related to neural activity. The relative paucity of noise related to motion and physiological cycles in the global signal suggests that for studies in anesthetized rodents, global signal regression is likely to do more harm than good, attenuating information about neural activity while making only modest reductions in other noise. For humans, where motion and physiological noise make greater contributions, the implications of the study are less clear, but at a minimum this work should motivate a cautious approach to the removal of the global signal, and an additional level of care in the interpretation of the results.

## ACKNOWLEDGEMENTS

This research was funded by the National Institutes of Health (1R01MH111416, 1R01NS078095, 1R01AG062581, 1R01EB029857S1, 1R01EB029857). Additional thanks to the Center for Systems Imaging Core (CSIC) at Emory University and the NIH/NIBIB T32 Computational Neural Engineering Training Program at Georgia Tech and Emory University.

## SUPPLEMENTARY MATERIALS

**Supplementary Table 1:**

|  | Length of isoflurane scan (minutes) | Time after disconnection of isoflurane that dexmedetomidine scan starts (minutes) | Length of dexmedetomidine scan (minutes) | Time after introduction of 0.5% isoflurane that isodex scan starts (minutes) | Length of isodex scan (minutes) |
| --- | --- | --- | --- | --- | --- |
| Subject 1 | 30 | 23 | 60 | 32 | 60 |
| Subject 2 | 30 | 19 | 60 | 7 | 60 |
| Subject 3 | 30 | 14 | 80 | 17 | 30 |
| Subject 4 | 30 | 8 | 80 | 23 | 20 |
| Subject 5 | 60 | 5 | 30 | 12 | 30 |
| Subject 6 | 40 | 17 | 60 | 14 | 40 |
| Subject 7 | 30 | 12 | 60 | 13 | 60 |
| Subject 8 | 30 | 13 | 60 | 18 | 90 |

**Supplementary Figure 1:**
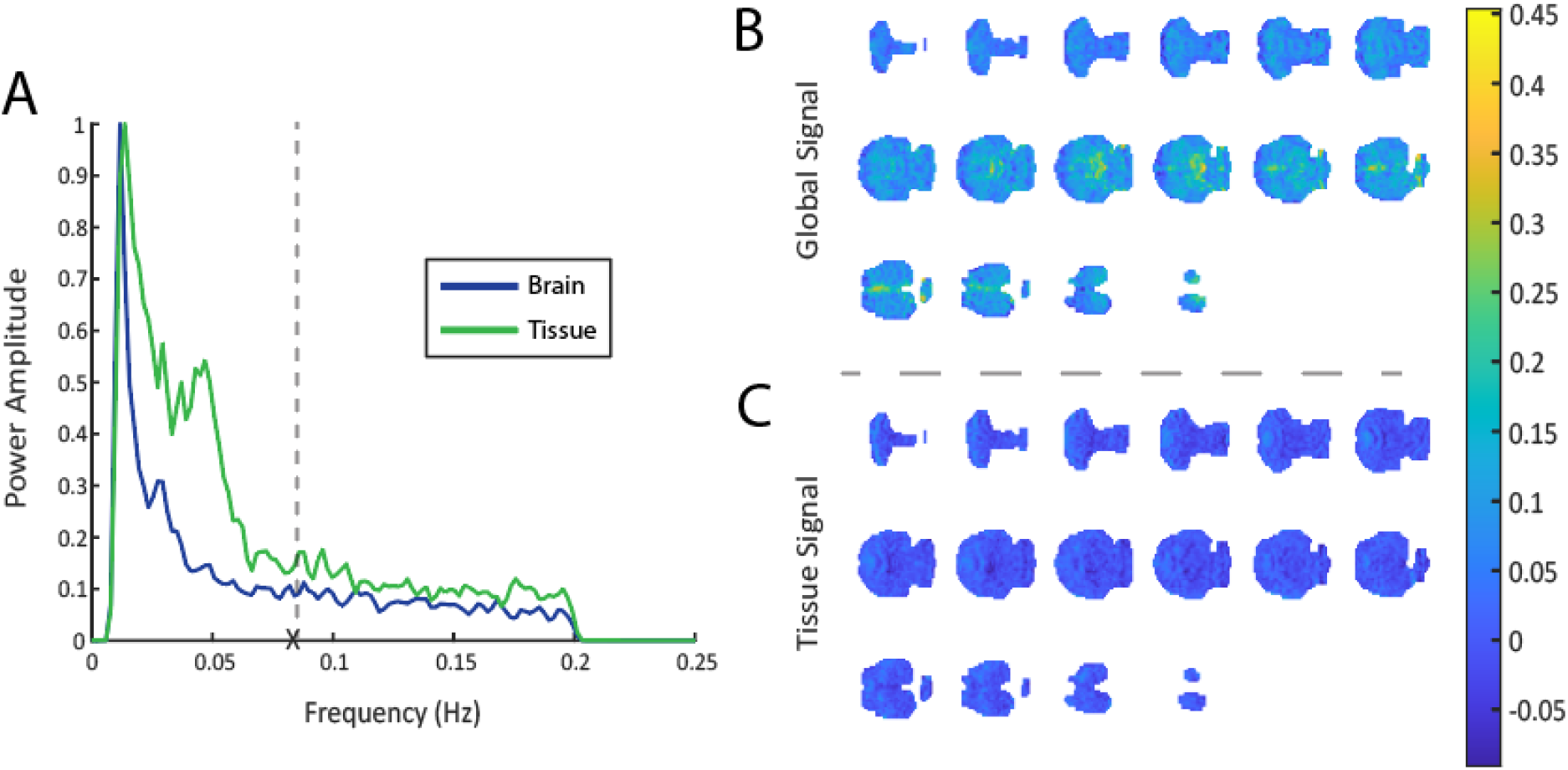
Analysis of the last 30 minutes of dexmedetomidine scans. To account for any lingering effects of isoflurane on the first 30 minutes of dexmedetomidine scans, we also looked at the PSD estimate and spatial correlation of the global and tissue signals in the last 30 minutes of these scans. For both analyses, we saw similar results to those presented in the main analysis. (A) The tissue signal consisted of a higher contribution from lower frequencies than the brain global signal, with the tissue signal having a fALFF of 0.5396 and the global signal having a fALFF of 0.5035. (B) A voxel-wise correlation to the global signal shows higher correlation to the global signal from medial brain structures, as was seen in the first 30 minutes. (C) The same voxel-wise analysis to the tissue signal shows lower overall correlation, as was seen in the first 30 minutes.

**Supplementary Figure 2:**
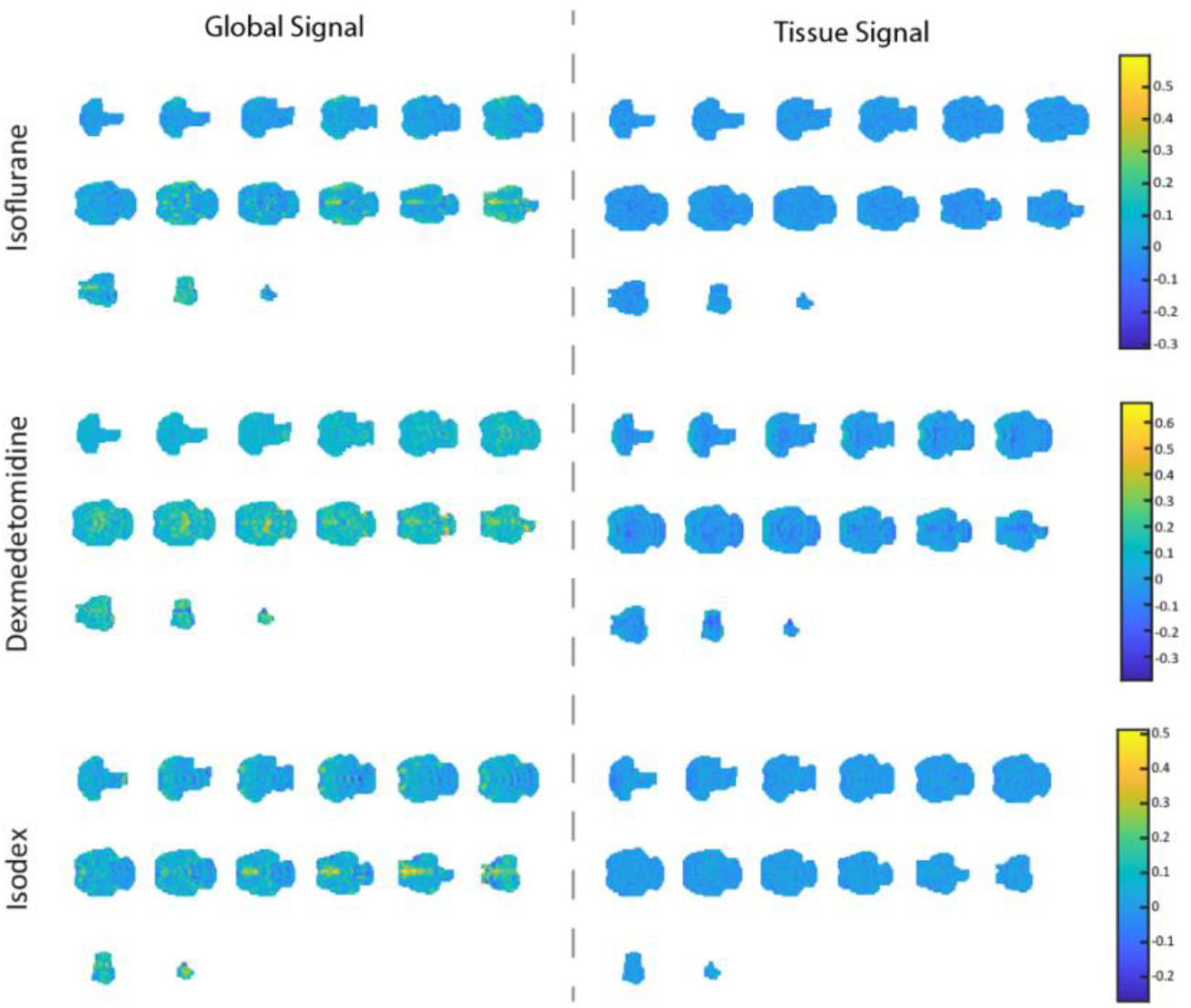
Correlation analysis in subject 1. This figure shows the individual voxel-wise correlation analysis to the global and tissue signals in subject 1.

**Supplementary Figure 3:**
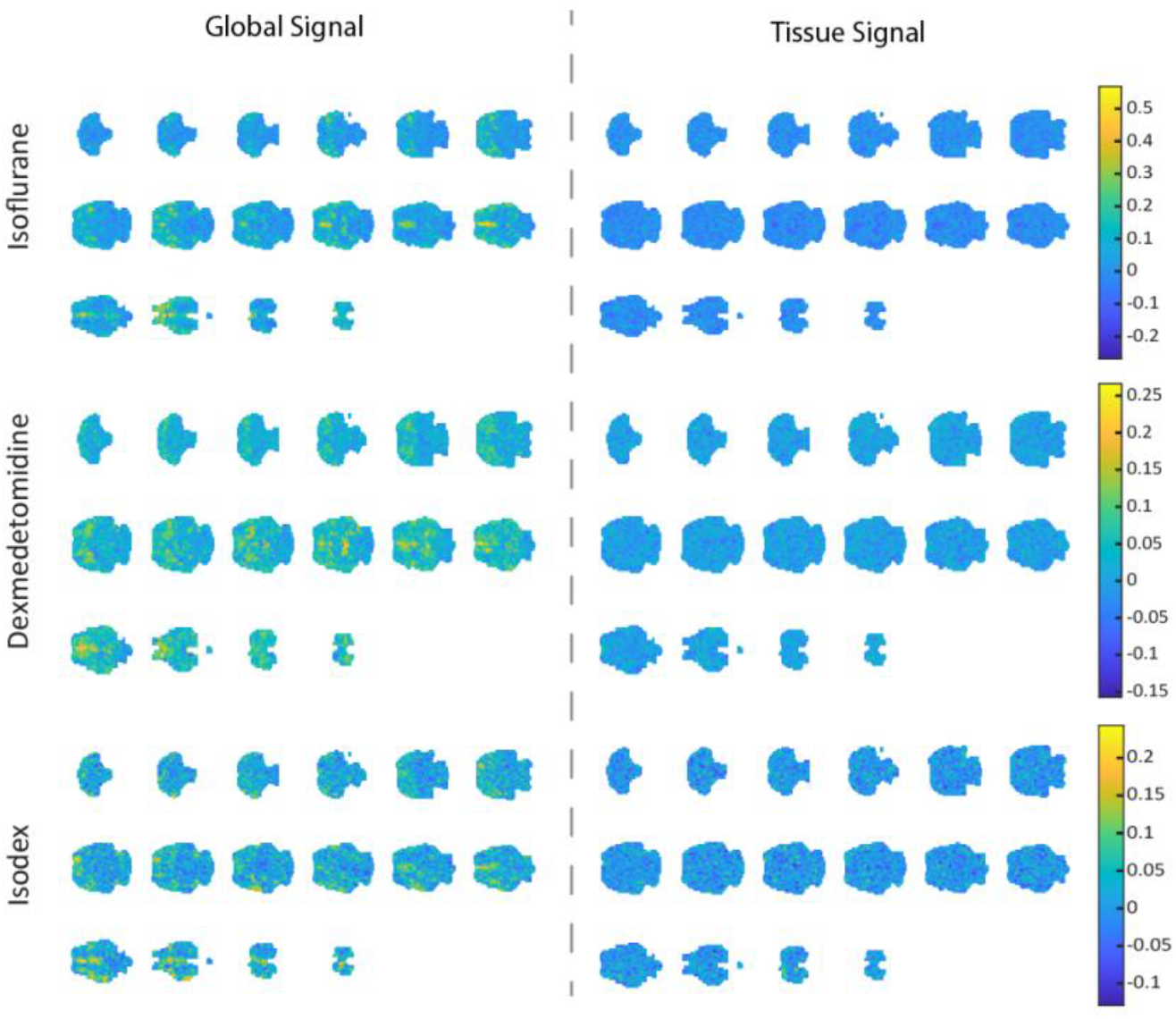
Correlation analysis in subject 2. This figure shows the individual voxel-wise correlation analysis to the global and tissue signals in subject 2.

**Supplementary Figure 4:**
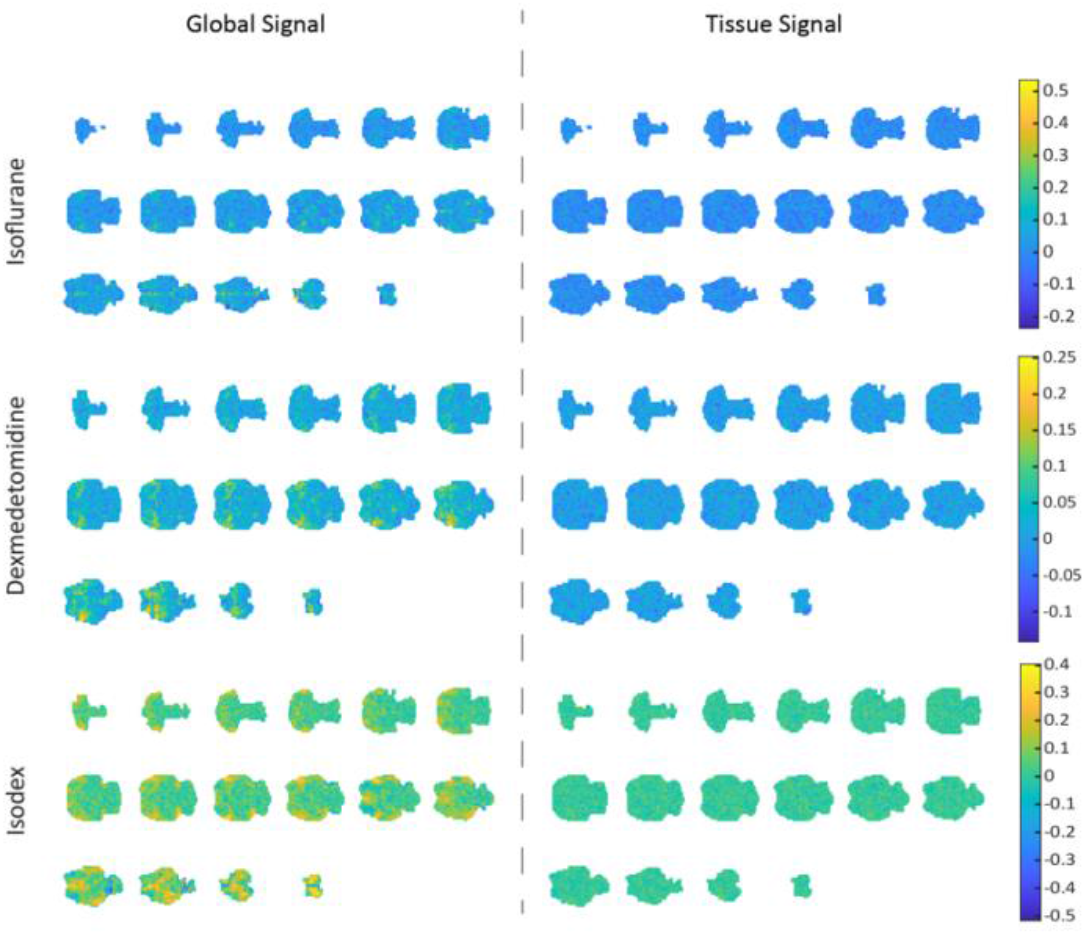
Correlation analysis in subject 3. This figure shows the individual voxel-wise correlation analysis to the global and tissue signals in subject 3.

**Supplementary Figure 5:**
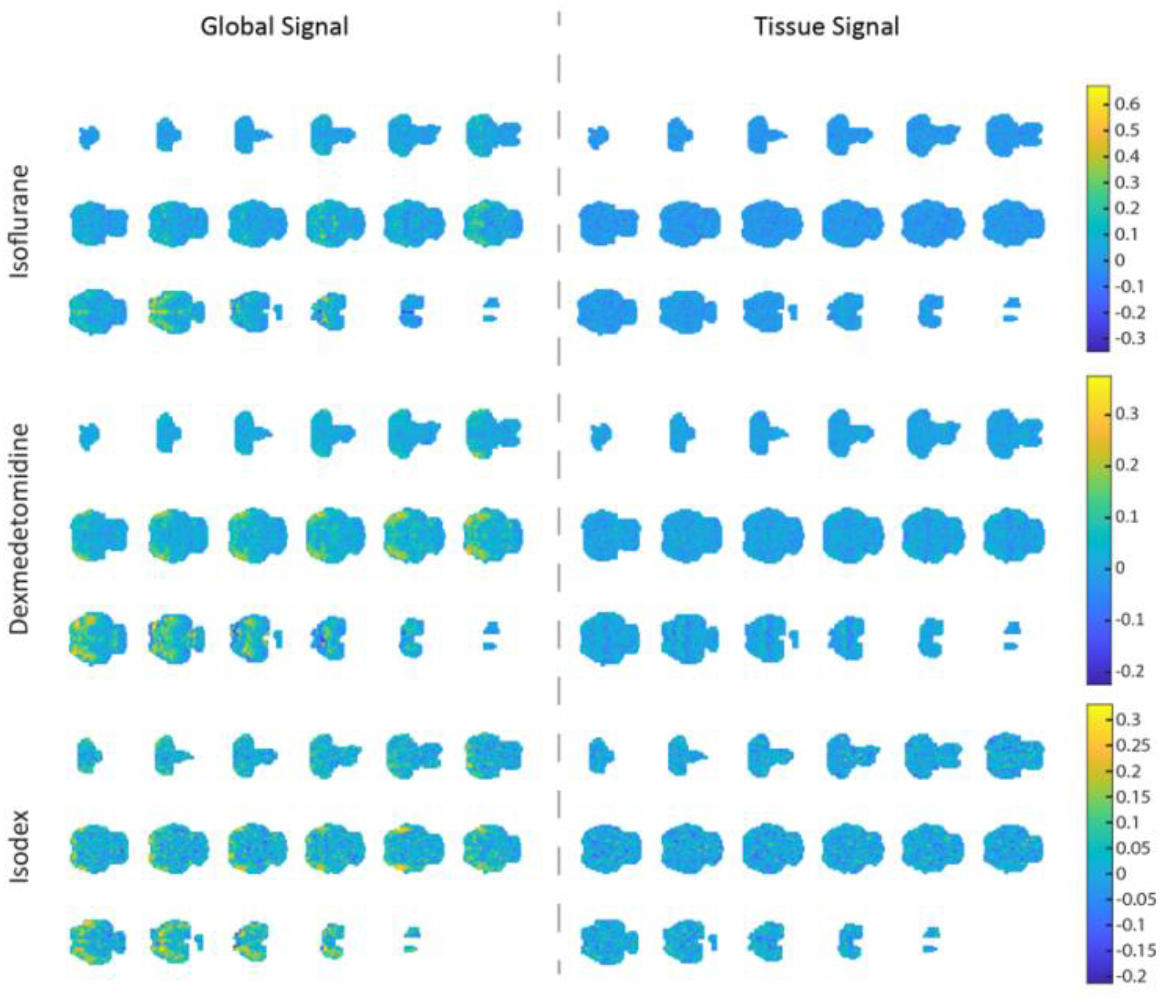
Correlation analysis in subject 4. This figure shows the individual voxel-wise correlation analysis to the global and tissue signals in subject 4.

**Supplementary Figure 6:**
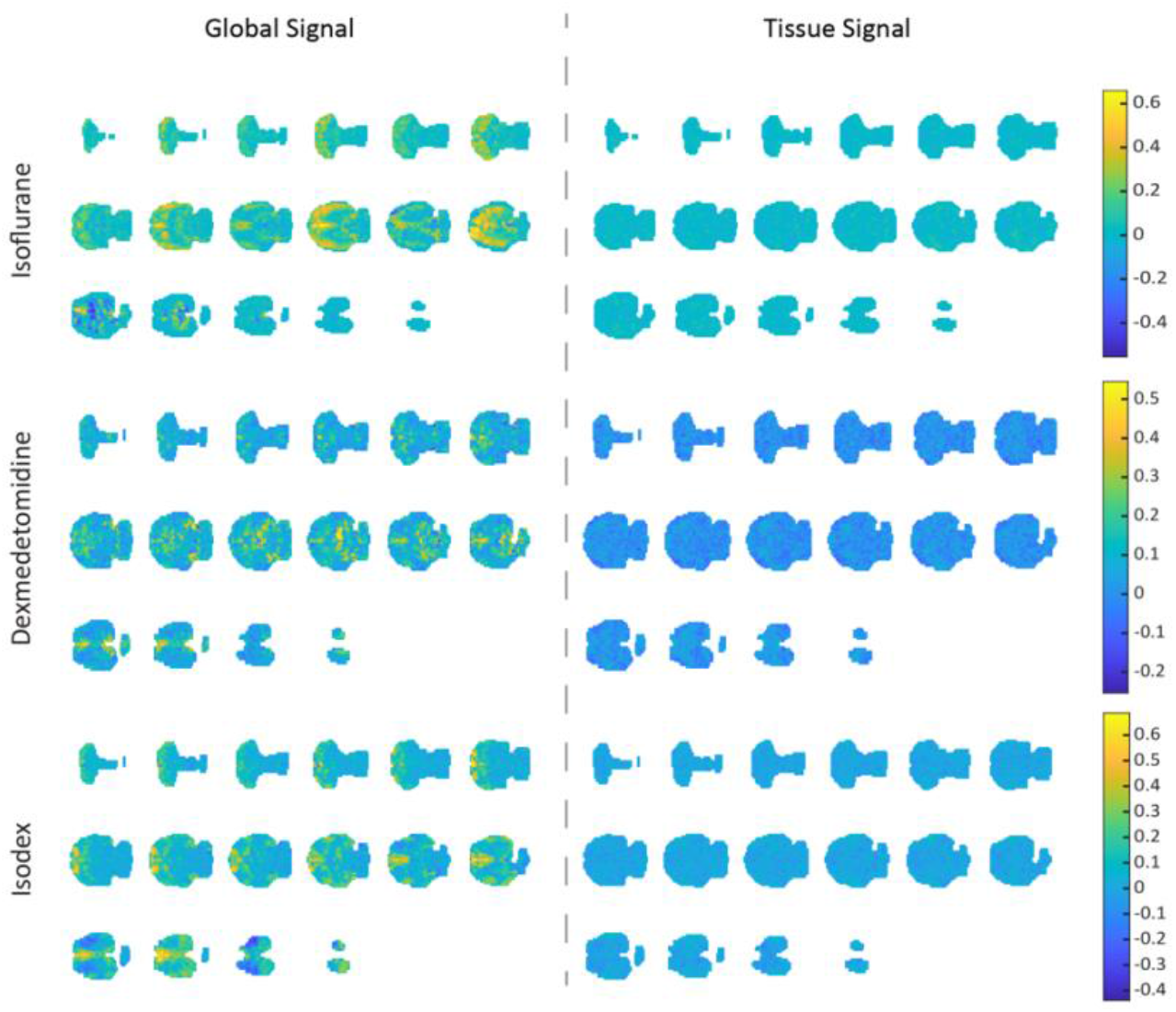
Correlation analysis in subject 5. This figure shows the individual voxel-wise correlation analysis to the global and tissue signals in subject 5.

**Supplementary Figure 7:**
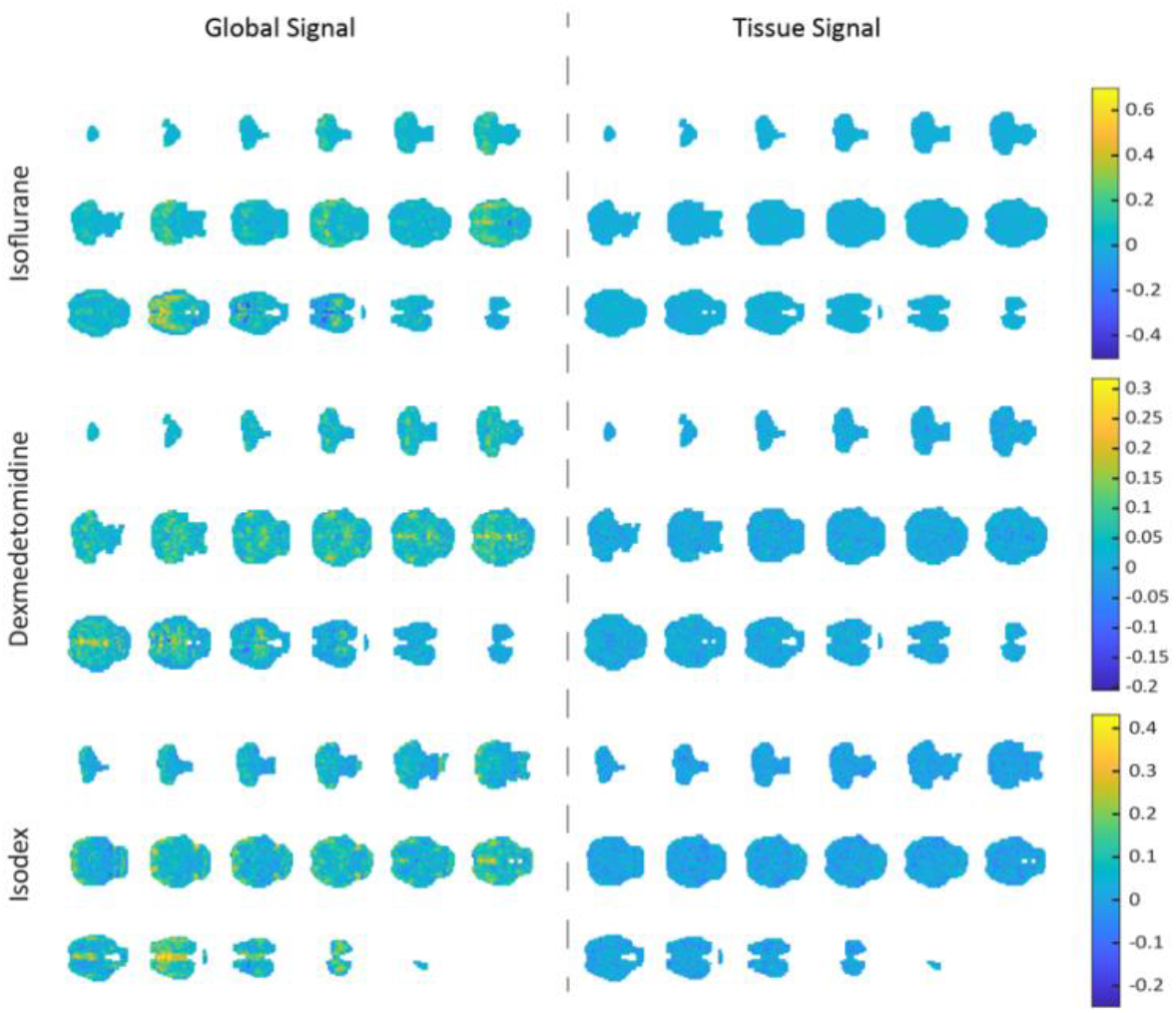
Correlation analysis in subject 6. This figure shows the individual voxel-wise correlation analysis to the global and tissue signals in subject 6.

**Supplementary Figure 8:**
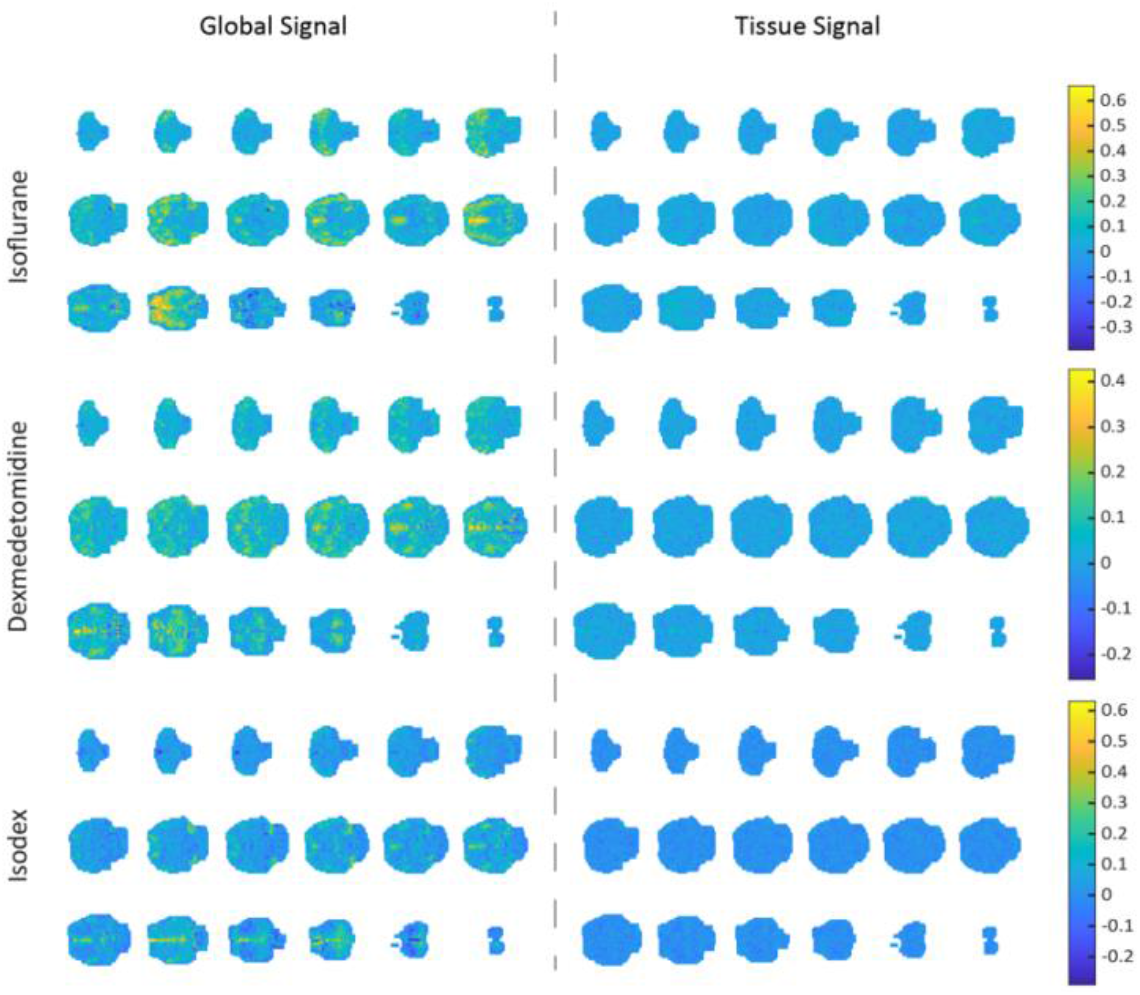
Correlation analysis in subject 7. This figure shows the individual voxel-wise correlation analysis to the global and tissue signals in subject 7.

**Supplementary Figure 9:**
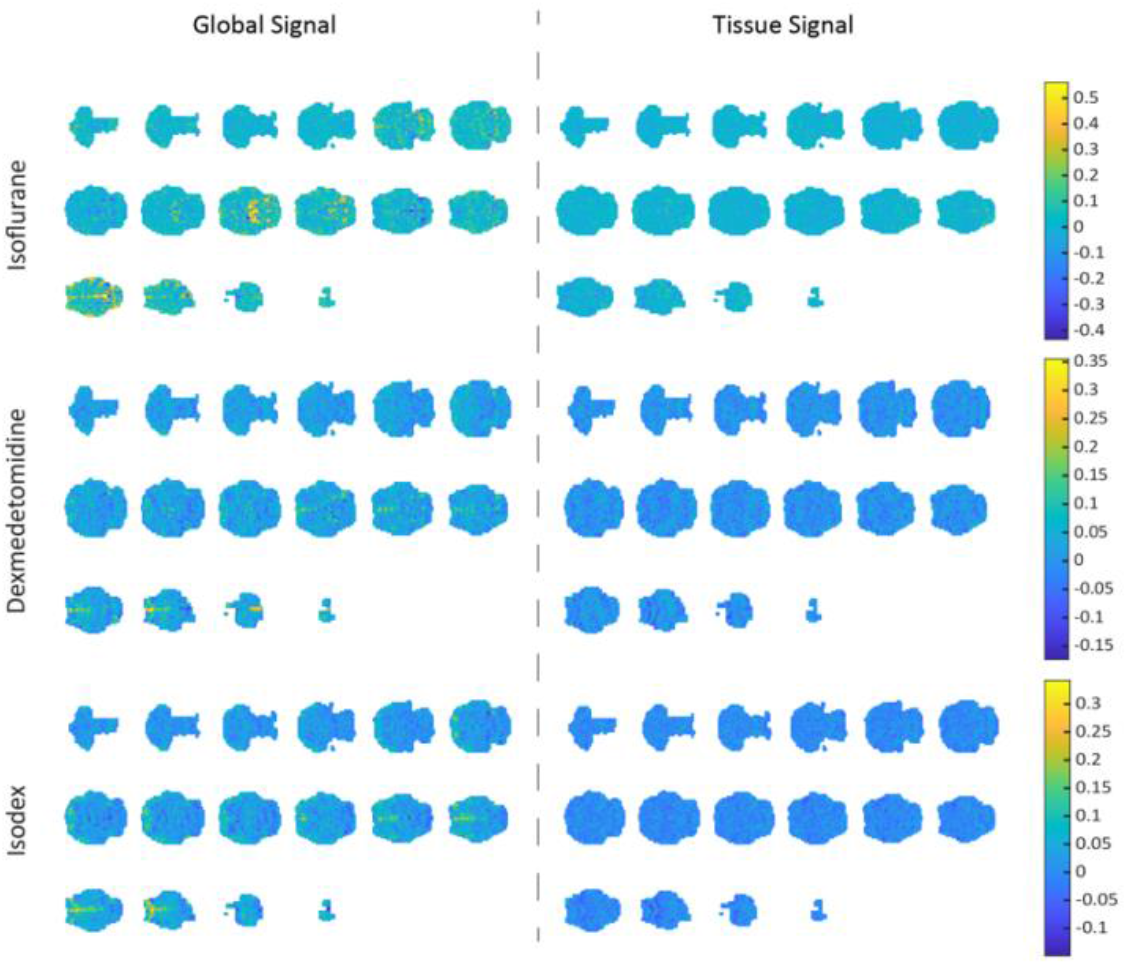
Correlation analysis in subject 8. This figure shows the individual voxel-wise correlation analysis to the global and tissue signals in subject 8.

